# Minoxidil hydrochloride impedes NLRP3 inflammasome activation via upregulation of AMPK-mediated autophagy

**DOI:** 10.64898/2026.04.06.716638

**Authors:** Sukhleen Kaur, Mehboob Ali, Anjum Shafeeq, Ziya Tufail, Lakhvinder Singh, Zabeer Ahmed, Ajay Kumar

## Abstract

NLRP3 inflammasome is a cytosolic multi-protein complex that plays a crucial role in the immune system, responding to various exogenous and endogenous stimuli by triggering protective inflammatory responses. However, aberrant NLRP3 inflammasome activation is implicated in numerous inflammatory diseases. Therefore, the NLRP3 inflammasome is an important pharmacological target for treating multiple diseases. In this context, we screened various US-FDA-approved drugs for NLRP3 inflammasome inhibition. We found that minoxidil hydrochloride (MXL) effectively inhibits the NLRP3 inflammasome, with IC50 values of 1.2 and 1.06 µM for IL-1β and IL- 18 release, respectively, thereby protecting J774A.1 cells from inflammatory cell death. Similarly, anti-NLRP3 activity was confirmed in bone marrow-derived macrophages (BMDMs), where MXL was found to prevent ASC oligomerization, thereby inhibiting the NLRP3 inflammasome and leading to CASP1 cleavage. Further investigation revealed that MXL also utilizes AMPK-mediated autophagy to modulate NLRP3 inflammasome activity. Using *siAMPK* and bafilomycin A1, an end-stage autophagy inhibitor, we elucidated crosstalk between the NLRP3 inflammasome and autophagic pathways, which was modulated by MXL. Furthermore, we demonstrated the efficacy of MXL in two different mouse models of inflammation involving the NLRP3 inflammasome. MXL at doses of 10 and 20 mg/kg effectively inhibited the activation of NLRP3 inflammasome by monosodium urate in the air pouch model and by ATP in the peritoneal inflammation model, as evidenced by reduced secretion of IL-1β and IL-18 in the lavage. Our study identifies MXL as a potent inhibitor of the NLRP3 inflammasome, warranting further investigation as a therapeutic agent for inflammatory diseases.

## 1 Introduction

Inflammation represents a critical component of the host immune defense mechanism, wherein diverse signaling pathways trigger the activation of the NLRP3 inflammasome in response to bacterial infections, pathogens, damaged mitochondria, damage-associated molecular patterns, and pathological states, among other stimuli (1–6). The NLRP3 inflammasome constitutes a fundamental component of the innate immune system, facilitating the maturation and release of pro-inflammatory cytokines, thereby initiating a cascade of inflammatory processes. Excessive activation of the NLRP3 inflammasome exacerbates the pathogenesis of numerous autoimmune, autoinflammatory, metabolic, cardiovascular, and neurodegenerative diseases (7–10). The heightened activation leads to an uncontrolled release of pro-inflammatory cytokines, perpetuating inflammatory responses and contributing to the progression and severity of these diseases. Hence, precise control of NLRP3 inflammasome is imperative, necessitating the augmentation of recycling pathways or cellular survival mechanisms, such as autophagy. Autophagy is a fundamental homeostatic process, ensuring cell viability by eliminating misfolded proteins, damaged organelles, and inflammasome activators through lysosomal degradation pathways. Recent investigations have garnered interest in pharmacologically inhibiting the NLRP3 inflammasome as a pivotal strategy for modulating inflammatory responses.

In this investigation, we explored the repurposing potential of minoxidil hydrochloride as a novel therapeutic agent. We found that minoxidil hydrochloride exhibits a robust inhibitory effect against the activation of the NLRP3 inflammasome induced by LPS and ATP in murine macrophages. Moreover, mechanistic investigations unveiled that minoxidil hydrochloride upregulates AMPK-mediated autophagy, thereby modulating the expression of proteins associated with inflammasome activation. Administration of minoxidil hydrochloride at two doses in two well-established in vivo inflammatory models demonstrated significant anti-NLRP3 inflammasome activity. This was evidenced by the marked reduction in the secretion of pro-inflammatory cytokines. These findings suggest that minoxidil hydrochloride has the potential to be repurposed as a therapeutic agent for the treatment of various inflammatory diseases, where the over-activation of the NLRP3 inflammasome exacerbates disease pathology.

## 2. Methods and Materials

### 2.1 Chemicals and Antibodies

Chemicals: DMEM (D1152), Phosphate buffer saline (D5652), Sodium bicarbonate (S5761), Penicillin (P3032), Streptomycin (S6501), HEPES (H3375), Lipopolysaccharide (L3129), Adenosine 5’-triphosphate (A6419), Bafilomycin A1 (B1793), DAPI (D9542), Paraformaldehyde (P6148), Triton X-100 (T8787), glycerol (G5516), Tween 20 (P7949), Trizma (T6066), SDS (L3771), Uric acid sodium salt (U2875) and Suberic acid (S1885) were purchased from Sigma-Aldrich. Acrylamide (193982), Phenylmethylsulfonyl (195381), Albumin Bovine Fraction V (160069) and Glycine (194825) were purchased from MP Biomedical. Fetal Bovine Serum (FBS) (10270106) and OPTI-MEM media (11058-021) were procured from Gibco. EDTA (Invitrogen-15575038), Skimmed milk (Himedia-GRM1254) and Strataclean resin (Agilent-400714-61) were used.

Antibodies: Anti-NLRP3 (15101S), pAMPK (2535S), MTOR (2972S), pMTOR (5536S), HRP-linked anti-rabbit IgG (7074S), HRP-linked anti-mouse IgG (7076S), anti-mouse IgG Alexa flour 488 (4408S), anti-rabbit IgG Alexa flour 488 (4412S) antibodies and siRNA AMPK (6620S) were purchased from Cell Signaling Technology (CST). All the CST antibodies were diluted (1:1000 times). Anti-ASC (SC-22514), CASP1 (SC-56036), BECN1 (SC-48341) and HRP-linked anti-goat IgG (SC-2354) antibodies were purchased from Santa Cruz Biotechnology. All the antibodies from Santicruz Biotechnology were diluted (1: 500 times), Anti-ACTB (A3854), working dilution (1:5000 times), anti-LC3B-II (L7543), working dilution (1:5000 times), and anti-p62/SQSTM1 (P0067), working dilution (1: 1000 times), were procured from Sigma-Aldrich. Anti-mIL-1β (AF-401-NA) working dilution (1:1000) was purchased from R&D Biotechnology.

Kits and reagents: Precision plus protein markers (161-0375) and Bradford reagent (Bio-Rad-500-0006) were purchased from Bio-Rad. ECL-kit (WBKLS0500) and PVDF Membrane (ISEQ00010) were obtained from Millipore. Mouse IL-1β ELISA kit (88-7013-88) and mouse IL-18 ELISA kit (88-50618-88) were purchased from Invitrogen. FuGENE HD (E2313) was purchased from Promega.

### 2.2 Cell culture

The J774A.1 murine macrophage cell line was procured from ECACC. These cells were cultured in DMEM media supplemented with 10% fetal bovine serum (FBS), penicillin (70 mg/L), streptomycin (100 mg/L), and NaHCO3 (3.7 g/L), in an environment maintained at 37°C with a constant supply of 5% CO2 and 95% humidity.

### 2.3 Bone Marrow-derived Macrophages

Balb/C mice were sacrificed and disinfected with 70% ethanol. Both femur and tibia were separated from the muscular layers and placed in PBS. To extract BMDMs, RPMI medium was used to flush each bone through a 25-gauge needle, after which the cells were gently dissociated by pipetting and passed through a cell strainer. The cells were differentiated for 7 days in RPMI medium supplemented with 10% heat-inactivated FBS and 30% L929 conditional medium.

### 2.4 Activation of NLRP3 inflammasome in BMDMs and J774A.1 macrophage cells

The activation of NLRP3 inflammasome in BMDMs and J774A.1 cells were induced by LPS and ATP. The cells were seeded in 24-well plates and cultured for 24 h. Following this incubation period, the cells were exposed to LPS (1 μg/ml) for 3.5 h in BMDMs and 5.5 h in J774A.1. Afterwards, the cells were treated with varying concentrations of MXL and standard inhibitor MCC 950 (100 nM), for 1 h in serum-free medium, and then stimulated with ATP (3 mM) for 30 min before the termination of the experiment.

### 2.5 Detection of cytokine levels through ELISA

After inducing the NLRP3 inflammasome and administering the specified drug treatments (as detailed earlier), supernatants were analyzed to measure the levels of pro-inflammatory cytokines IL-1β and IL-18 using ELISA, following the guidelines provided by the manufacturer (Invitrogen). Cytokine levels were normalized by dividing the measured values by the total protein content in each sample.

### 2.6 Lactate dehydrogenase activity (LDH)

LDH was measured in the supernatants of J774A.1 cells using the lactate dehydrogenase assay kit according to the manufacturer’s instructions (Sigma-Aldrich).

### 2.7 Immunoblotting

The levels of mature IL-1β and caspase-1 were assessed in the supernatant. The supernatant was incubated with strataclean resin at 4°C for 1 h under continuous rotation to ensure efficient protein binding to the resin. Following a 10-minute centrifugation, 2X Laemmli dye was mixed with the resin pellet, and samples were heated at 95°C for 10 minutes. The prepared lysates were then subjected to protein separation via SDS-PAGE. For cellular lysates, cells were lysed in RIPA buffer containing PMSF (2 mM), Na3VO4 (0.5 mM), NaF (50 mM), and 1% protease inhibitor cocktail, followed by centrifugation at 12,000 rpm for 20 min at 4°C (11). The cellular lysates were applied to SDS-PAGE for protein separation, followed by transfer to a PVDF membrane for 2 hours at 4°C. The membranes were then blocked with 5% BSA or 5% skimmed milk for 1 hour at room temperature, followed by overnight incubation with primary antibody at 4°C. Subsequently, the membranes were incubated with an HRP-conjugated secondary antibody for 2 hours at room temperature. Protein bands were detected using a chemiluminescent HRP substrate (Millipore) and visualized with the Chemidoc system (Syngene Gchemi XT4).

### 2.8 Oligomerization of ASC

Following drug treatments under conditions activating the NLRP3 inflammasome, cells were lysed in a cold buffer consisting of 1% protease inhibitor cocktail, 1 mM sodium orthovanadate, 150 mM KCl, 0.1 mM PMSF, 20 mM HEPES-KOH, and 1% NP-40. The lysed cells were centrifugated at 330g for 10 min at 4°C, and the supernatant was collected for subsequent protein expression analysis via immunoblotting. Cellular pellets were washed with PBS before being resuspended in 500 μL of cold PBS. Subsequently, a 2 mM concentration of suberic acid was added to the pellets, which were then incubated for 30 minutes at 37°C to facilitate cross-linking of ASC proteins (11). Following incubation, the cell pellets were centrifuged at 330 g for 10 minutes at 4 °C. After centrifugation, 2X Laemmli dye was added to the cell pellets. The resulting cell lysates were heated at 95°C for 10 minutes, after which protein analysis was performed by western blotting.

### 2.9 Flow Cytometry

Propidium iodide staining was used to measure cellular death. After inducing the NLRP3 inflammasome and administering the specified drug treatments (as detailed earlier, PI was added to a final concentration of 1 µg/ml and incubated in the dark for 15 min prior to termination. Cells were then washed and collected in PBS for the analysis of PI fluorescence using flow cytometry.

### 2.10 Confocal microscopy

For confocal microscopic examination of protein localization, cells were cultured in a 6-well plate over coverslips. Following various treatments, cells underwent triple PBS washes and fixation in 4% paraformaldehyde for 30 minutes. Permeabilization was achieved using 0.2% Triton X-100 for 7 minutes, followed by blocking with buffer containing 2% BSA and 0.2% Triton X-100 for 1 h. Subsequently, cells were incubated overnight with the primary antibody at 4°C. The following day, cells were treated with secondary antibody (Alexa flour 488) for 1 h at room temperature and underwent triple PBS washes. Nuclei were counterstained with DAPI, and coverslips were mounted with mounting medium. Images were captured using Yokogawa CQ1 Benchtop High-Content Analysis System at 40x or 60x magnification.

### 2.11 Transfection of cells with AMPK siRNA

For AMPK transfection, cells were incubated in OPTI-MEM medium and transfected with AMPK siRNA using FuGENE HD reagent for 24 hours. Following transfection, the cells were exposed to LPS for 5.5 h, subsequently, the cells were treated with 5 and 10 μM MXL under conditions that activate the NLRP3 inflammasome. The samples were then analyzed by western blotting.

### 2.12 Drug Formulation

For in vivo studies, MXL was prepared in a solution comprising 5% DMSO, 30% PEG 400, and 20% PEG 200 in distilled water. Both LPS and ATP were formulated in PBS.

### 2.13 LPS and ATP-induced model of peritoneal inflammation

For model development, female Balb/c mice were divided into five groups: (1) Control, (2) LPS, (3) LPS+ATP, (4) LPS+ATP+MXL (10 mg/kg), and (5) LPS+ATP+MXL (20 mg/kg), with each group containing five animals (n=5). Thirty minutes prior to the LPS treatment (4h incubation), mice were administered intraperitoneally (i.p.) with two different MXL doses (10 and 20 mg/kg). Following LPS treatment incubation, the mice received an intraperitoneal injection of ATP (100 mM) for 15 minutes. Post-treatment, the animals were sacrificed, and 3 mL of incomplete DMEM medium was administered intraperitoneally. The peritoneal lavage fluid was then collected along with the medium and analyzed for pro-inflammatory cytokine levels using ELISA.

### 2.14 Air Pouch Model

The air pouch model was developed to investigate the effects of MXL on monosodium urate (MSU) crystal-induced inflammation. Balb/c mice were randomly allocated into five groups, each containing five animals (n=5): (1) Control, (2) MSU, (3) Colchicine, (4) MXL (10 mg/kg), and (5) MXL (20 mg/kg). To induce the model, 4 mL of sterile air was injected into the dorsal subcutaneous space of the mice on the first day, with a repeat injection on the third day to maintain the air pouch. On the sixth day, mice were treated with MXL (10 mg/kg and 20 mg/kg) and colchicine. Thirty minutes after the drug treatment, MSU (3 mg/ml) was injected into the induced air pouches. Six hours post-MSU treatment, the mice were sacrificed, and air pouch lavages were collected after injection of incomplete DMEM. The pro-inflammatory cytokine levels in the collected lavages were measured using ELISA.

### 2.15 Animals and Ethical Clearance

The Balb/c mice were maintained under a 12-hour light/dark cycle in a controlled environment with a temperature ranging from 65–75°F (approximately 18-23°C) and humidity levels between 40–60%. The mice had free access to food and water (ad libitum). Prior to the study, all animals were acclimated for one week under standard laboratory conditions and were drug-naive, with no prior procedures having been performed. The experimental protocols were approved by the Institutional Animal Ethics Committee (IAEC) (IAEC approval number-305/81/8/2022) and adhered to the guidelines set by the Committee for Control and Supervision of Experiments on Animals (CCSEA), Ministry of Environment and Forest, Government of India.

### 2.16 Statistical analysis

The data are presented as mean ± standard deviation (SD). Statistical analysis was performed using GraphPad Prism software (version 9). One-way analysis of variance (ANOVA) followed by a Bonferroni post hoc test was conducted at a 95% confidence level. Formal normality testing (e.g., Shapiro-Wilk) was intentionally omitted due to its well-known lack of statistical power at n = 3; parametric assumptions were instead based on established historical distribution patterns for the assays used in this study and comparable group variances. A p-value of less than 0.05 was considered statistically significant. Significance levels were denoted as follows: **** p<0.0001; ***p<0.001; **p<0.01; *p<0.05.

## 3. Results

### 3.1 Screening of US FDA-approved drugs for NLRP3 inflammasome activation

Some US FDA-approved drugs were randomly selected to assess their anti-inflammatory activity against the activated NLRP3 inflammasome. Among these, Minoxidil (MXL) and Fluvoxamine were found to be significant inhibitors of the NLRP3 inflammasome, showing more than 40% reduction in the release of IL-1β and IL-18 (Figures 1A and B). Our research group previously demonstrated that fluvoxamine exhibits a potent anti-NLRP3 activity both in vitro and in vivo in the 5XFAD transgenic mice (11). Therefore, this study was focused on investigating the anti-NLRP3 inflammasome activity of MXL. After confirming the inhibitory effect of MXL on the NLRP3 inflammasome, we calculated the IC50 value of MXL in LPS-primed and ATP-activated J774A.1 cells after various treatments with MXL (0.039 µM to 20 µM). We measured the levels of IL-1β and IL-18 secreted into the media by J774A.1 cells. The IC50 values for the inflammatory cytokines IL-1β and IL-18 were calculated to be 1.2 and 1.06 µM, respectively (Figure 1C-F).

**Figure 1:**
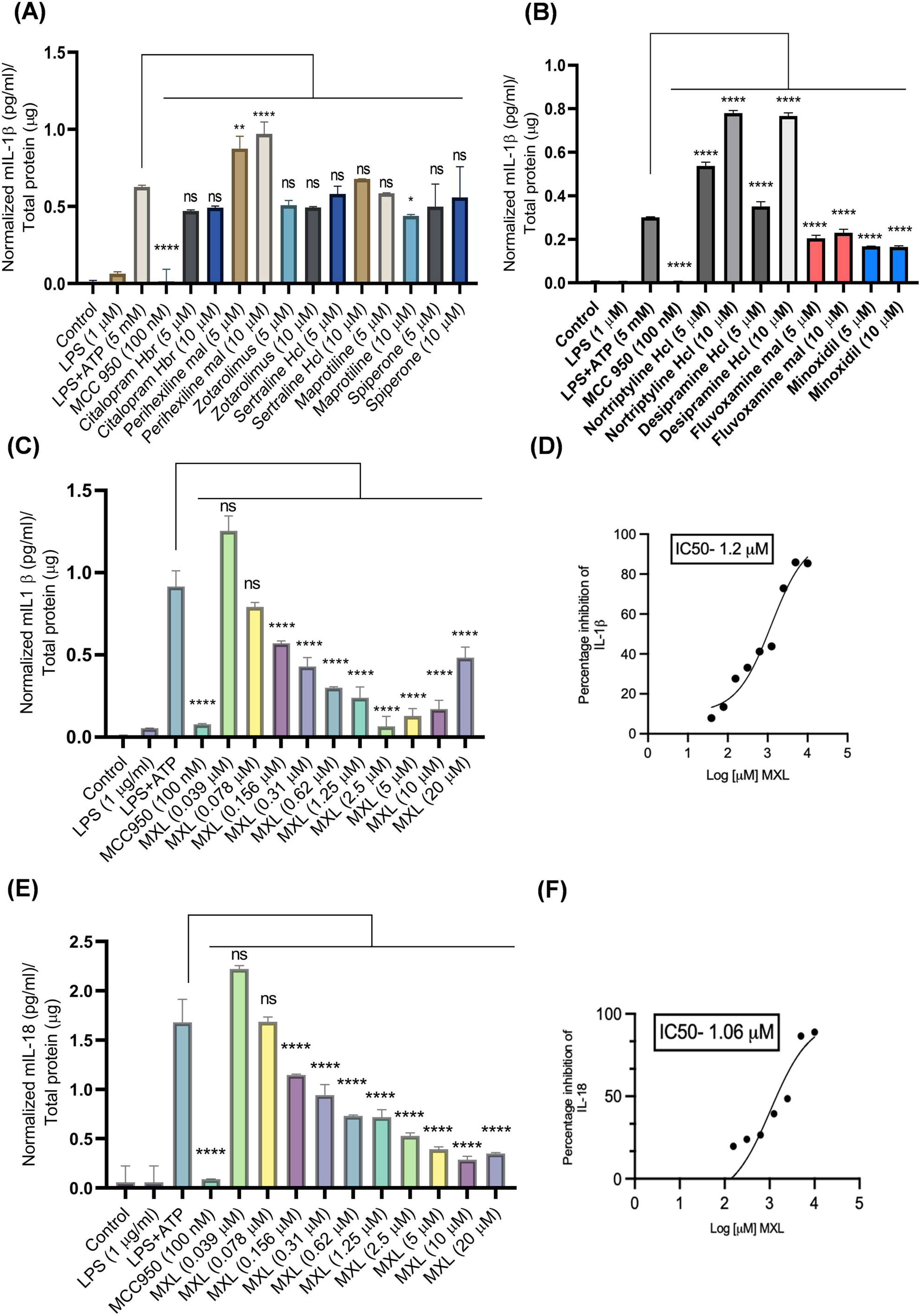
Screening of FDA-approved drugs for NLRP3 inflammasome activation. J774A.1 cells were primed with LPS and subsequently activated with ATP. The concentrations of the released pro-inflammatory cytokines IL-1β and IL-18 were quantified using ELISA. Normalized IL-1β levels as depicted in the graphs **(A)** and **(B)**. **(C)** Graph showing the normalized levels of IL-1β and **(D)** IC50 value of MXL for IL-1β in J774A.1 cells. **(E)** Graph illustrating the normalized levels of IL-18. **(F)** IC50 value of MXL for IL-18. Results are expressed as Mean±SD from three independent experiments (n=3). Data were analyzed using one-way ANOVA followed by Bonferroni correction, considering *p< 0.05 statistically significant ( ****p < 0.0001, ***p < 0.001, **p< 0.01 and ns = not significant).

### 3.2 MXL inhibited the activation of the NLRP3 inflammasome in bone marrow-derived macrophages

After identifying minoxidil as a potential inhibitor of the NLRP3 inflammasome in J774A.1 cells, we confirmed MXL’s anti-NLRP3 activity in bone marrow-derived macrophages (BMDMs). We found that MXL effectively inhibited the LPS+ATP-induced activation of the NLRP3 inflammasome with an IC50 value of 2.6 µM for IL-1β and 2.8 µM for IL-18 (Figures 2A-D). We further assessed the toxicity of MXL in J774A.1 and BMDM cells to determine whether the anti-NLRP3 activity is attributable to MXL toxicity. We found that both J774A.1 and BMDM cells treated with MXL did not exhibit significant toxicity at concentrations up to 100 µM, even after 24 h of treatment (Figures 2E and F). The IC50 values for MXL in both cell types were >100 µM.

**Figure 2:**
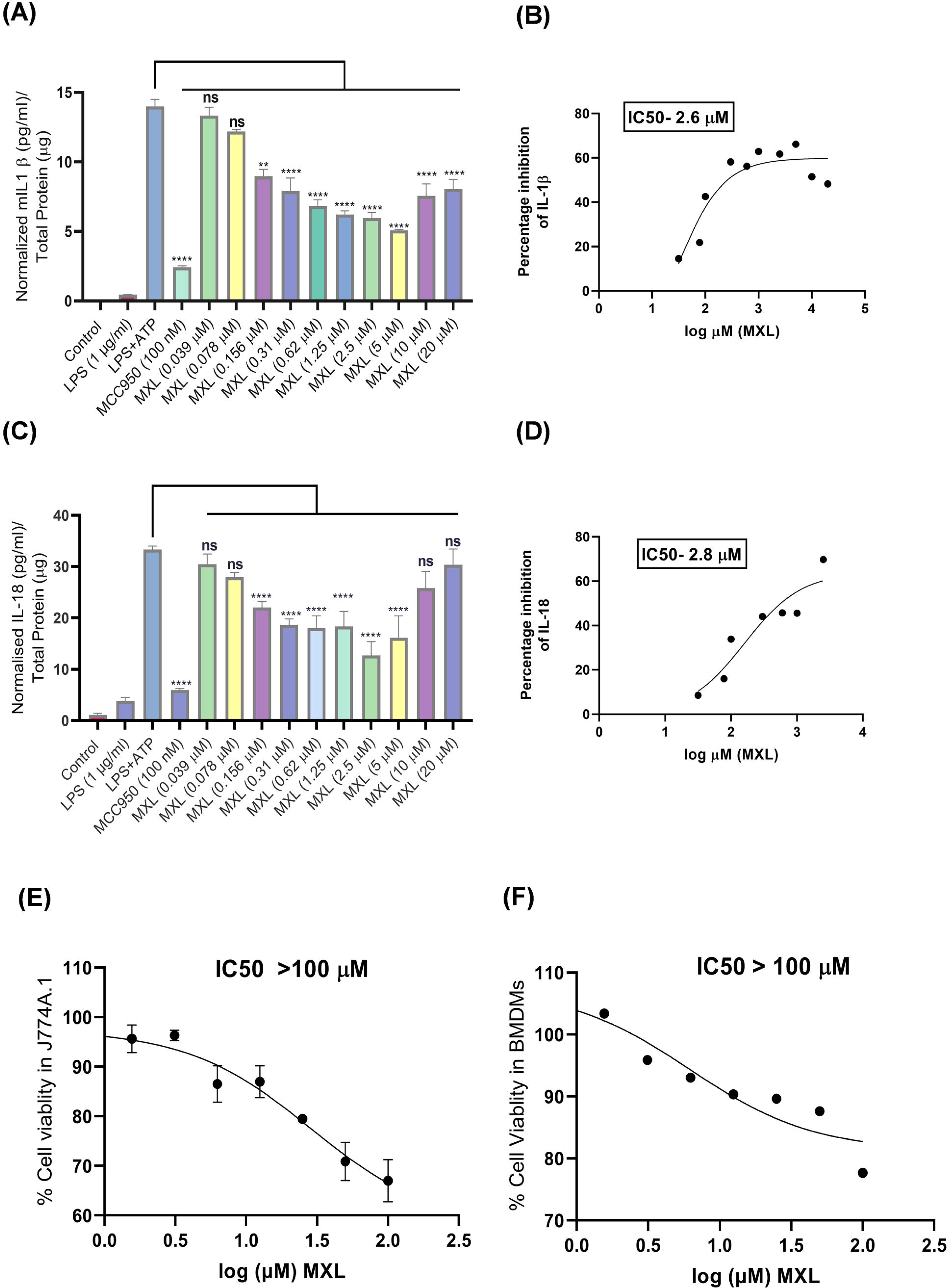
Effect of MXL on inhibition of NLRP3 inflammasome in BMDMs. **(A-B)** Concentration-dependent effect of MXL on the inhibition of ATP-mediated induction of NLRP3 inflammasome and subsequent release of IL-1β and **(C-D)** IL-18. **(E-F)** MTT assay to assess the toxicity of MXL in J774A.1 and BMDMs after 24 h of treatment. The results are expressed as mean ± SD from three independent experiments (n=3). Data were analyzed using one-way ANOVA followed by Bonferroni correction, considering *p< 0.05 statistically significant (****p < 0.0001, ***p < 0.001, **p< 0.01, and ns = not significant).

### 3.3 MXL inhibited the activation of the NLRP3 inflammasome to mitigate inflammatory cell death in J774A.1 cells

After a preliminary assessment of MXL’s effects on NLRP3 inflammasome activity, we selected 5 and 10 µM MXL concentrations for further studies based on its efficacy against both IL-18 and IL-1β. We further sought to elucidate the mechanism underlying its activity by priming J774A.1 cells with LPS and treating them with MXL for 1 h before ATP stimulation. The Western blot analysis indicated that in the presence of LPS and ATP, release of cleaved IL-1β and cleaved caspase-1 was significantly inhibited by MXL treatment in comparison to LPS+ATP-treated cells (Figures 3A-C). However, the expression of pro-IL-1β remained unaltered after MXL treatment at 5 and 10 µM. Intriguingly, MXL significantly reduced NLRP3 protein levels under similar treatment conditions (Figure 3A). After gaining an initial understanding of the mechanism of action of MXL in inhibiting the NLRP3 inflammasome, we sought to determine whether MXL treatment could rescue cells from death. Therefore, we used propidium iodide (PI) staining to assess cell death under inflammasome-activating conditions, with or without MXL. In the flow cytometric analysis, we found that after 5.5 hours of LPS treatment, 39% of the cells stained positive for PI; this number increased to 53% after adding ATP for 30 minutes (Figures 3E and F). Whereas the number of PI-stained cells was reduced to 15% when treated with 5 µM MXL and to 24% when treated with 10 µM MXL. The number of PI-stained cells was reduced to 15% with the standard drug MCC950 (Figures 3E and F). We also confirmed the protective effect of MXL by the LDH release assay in J774A.1 cells. We found that LDH release was reduced to 39% with 5 µM MXL and to 49% with 10 µM MXL. Whereas it was reduced to 25% for cells treated with the reference standard MCC950 (Figure 3F).

**Figure 3:**
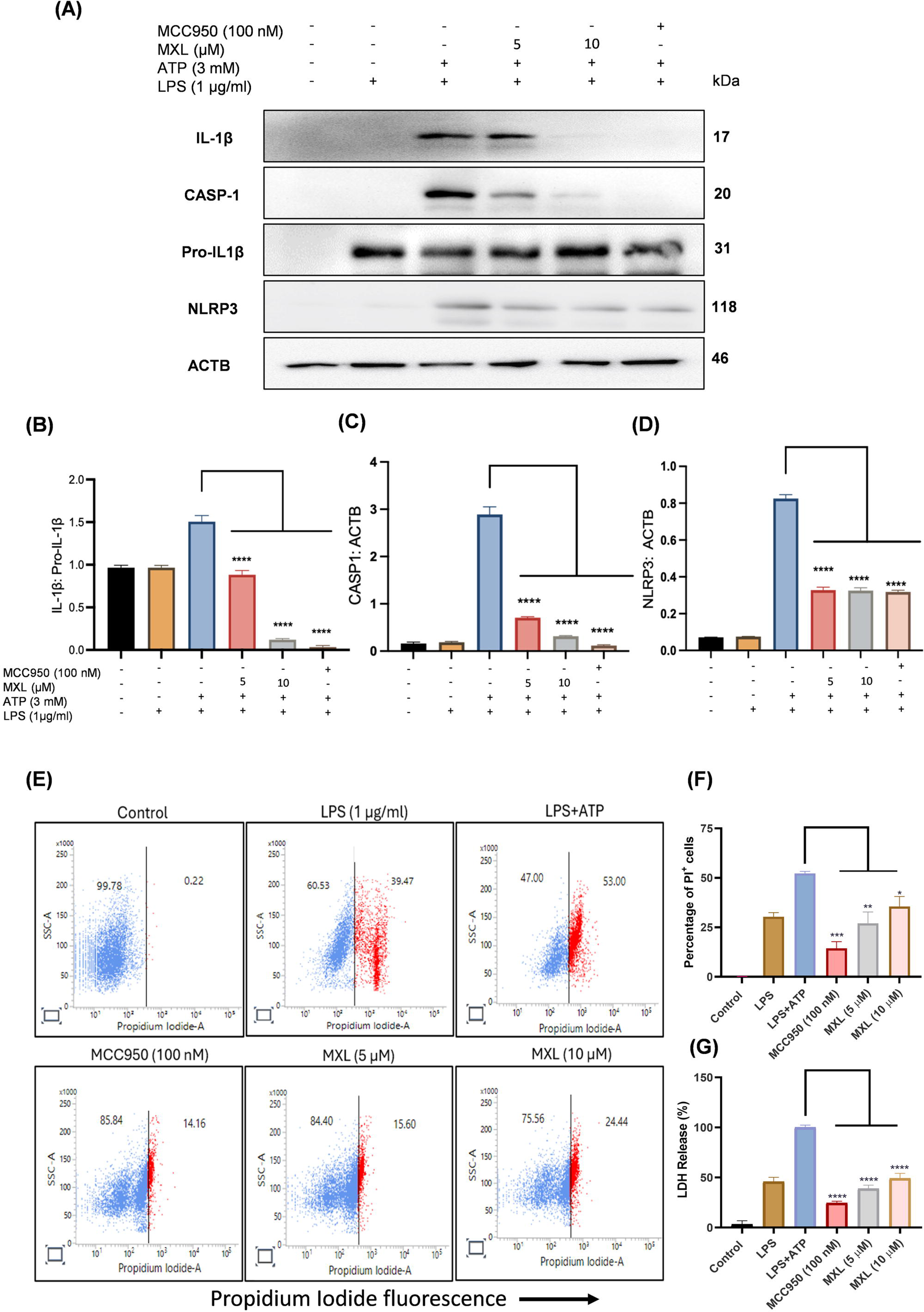
MXL inhibited the activation of NLRP3 inflammasome-mediated cell death. **(A)** Immunoblot analysis demonstrating the suppression of the NLRP3 inflammasome complex by MXL, including IL-1β, CASP1 and NLRP3. Densitometric measurements of immunoblots **(B)** IL-1β: Pro IL-1β, **(C)** CASP1: ACTB and **(D)** NLRP3: ACTB. **(E)** Representative dot plot of (PI) fluorescence analyzed by flow cytometry. **(F)** Quantification of the representative flow cytometry dot plots showing PI fluorescence. **(G)** LDH released in the supernatants of J774A.1 cells was measured by the LDH activity assay kit. The data are presented as the mean ± SD derived from the three independent experiments (n=3). Statistical analysis was conducted using one-way ANOVA, followed by a post-hoc Bonferroni test. A p-value of less than 0.05 was considered statistically significant, with the following significance levels: ****p < 0.0001, ***p < 0.001, **p< 0.01, *p< 0.05 and ns = not significant.

### 3.4 MXL reduced ASC oligomerization and speck formation

Under activated inflammasome conditions, adapter ASC proteins form a large structure called an ASC oligomer. So, we further explored the effect of MXL on ASC oligomerization by western blotting. J774A.1 cells treated with MXL showed significant inhibition of ASC oligomerization, dimerization, and monomer formation when compared to LPS+ATP-treated cells (Figure 4A and B). These results were also validated by using confocal microscopy, where MXL treatment significantly reduced the number of ASC speck formations compared to LPS+ATP-treated cells (Figure 4C and Supplementary Figure S1)

**Figure 4:**
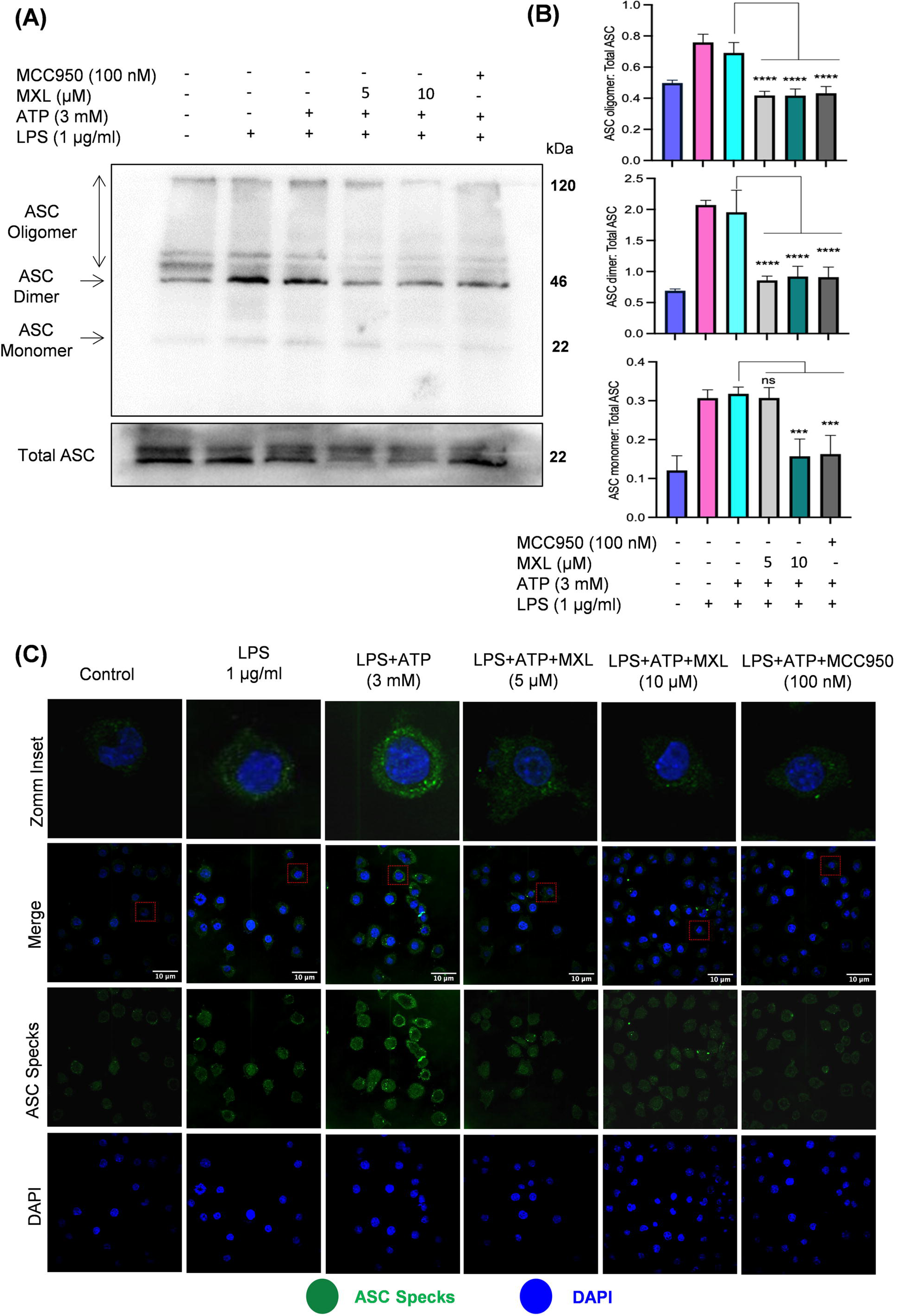
MXL reduced the ASC oligomerization and speck formation Following the activation of the NLRP3 inflammasome, the cross-linking agent suberic acid was employed to induce the oligomerization of ASC. This process facilitates the formation of ASC complexes, which are essential for inflammasome assembly and function. **(A)** Immunoblots analysis depicting ASC oligomerization. (**B)** Densitometric analysis of immunoblots revealed significant changes in ASC oligomerization, presented as ratio of ASC oligomer, ASC dimer, and ASC monomer to total ASC. **(C)** Representative confocal microscopy images illustrating ASC speck formation in J774A.1 cells, typically observed following NLRP3 inflammasome activation. The data represent the mean ± SD from three independent experiments (n=3), and the statistical comparison of the samples was performed by a one-way ANOVA, followed by the Bonferroni post-hoc test to account for multiple comparisons. A p-value of less than 0.05 was regarded as statistically significant. The levels of significance were indicated as follows: ****p < 0.0001, ***p < 0.001, **p< 0.01, *p< 0.05 and ns = not significant.

### 3.5 MXL induced autophagy via the AMPK pathway under inflammasome activation conditions in J774A.1 cells

Autophagy is an important regulatory mechanism of NLRP3 inflammasome activity. Therefore, we decided to explore its involvement in MXL-mediated inhibition of NLRP3 inflammasome. AMPK is the key enzyme involved in the induction of autophagy; therefore, we analyzed the AMPK activation by MXL under inflammasome-activating conditions. We found that MXL treatment significantly increased the expression of pAMPK (Thr172) compared with untreated controls and LPS/ATP-treated cells. Furthermore, the downstream proteins pmTOR (Ser2448) and 4E-BP1 were significantly inhibited by MXL (Figures 5A and 5 B). Further, the expression of the autophagic proteins BECN1 and LC3B-II was significantly upregulated by MXL treatment, whereas SQSTM1 expression was significantly downregulated (Figures 5A and B).

**Figure 5:**
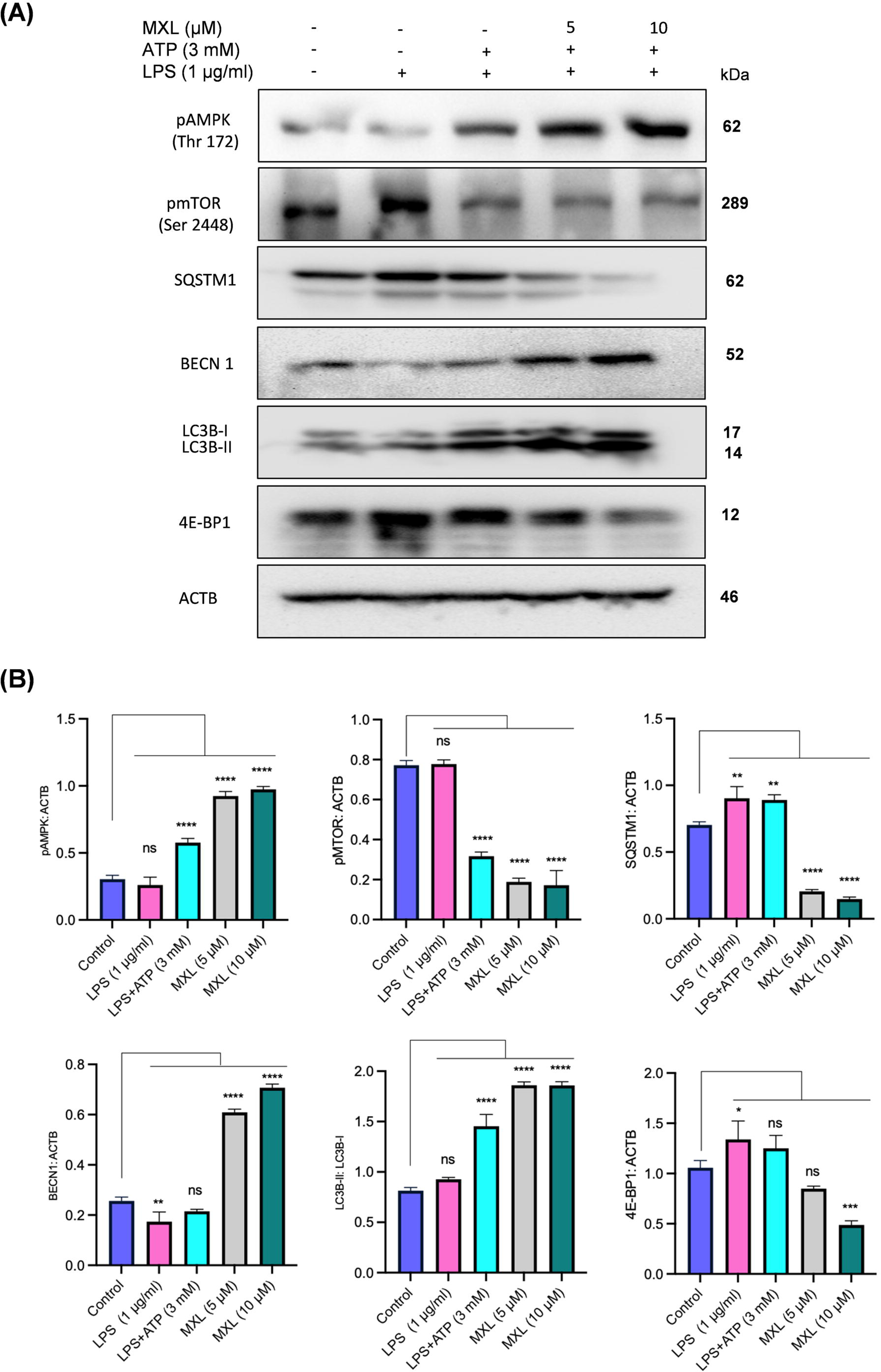
MXL induced autophagy via the AMPK pathway under the inflammasome activation condition in J774A.1 cells After activating the NLRP3 inflammasome, cell lysates were subjected to western blot analysis to assess the expression levels of autophagic proteins. **(A)** Immunoblots of autophagic proteins and, **(B)** Densitometric quantification of pAMPK: ACTB, pMTOR: ACTB, SQSTM1: ACTB, BECN1: ACTB, LC3B-II: LC3B-I, 4E-BP1: ACTB. The data represent the mean ± SD from three independent experiments (n=3). Statistical analysis was conducted using one-way ANOVA, followed by the Bonferroni post-hoc test. A p-value <0.05 was deemed statistically significant, with significance levels denoted as ****p < 0.0001, ***p < 0.001, **p< 0.01, *p< 0.05, and ns indicating not significant.

### 3.6 Pharmacological and genetic repression of autophagy limited the inhibitory effect of MXL on LPS and ATP-induced NLRP3 inflammasome activation

To investigate the correlation between MXL-induced autophagy and suppression of NLRP3 inflammasome, we used bafilomycin A1 (baf. A1) as a pharmacological inhibitor of autophagy. LPS/ATP-activated J774A.1 cells were treated with MXL in the presence or absence of Baf. A1 for 1 h. We found a clear inhibition of NLRP3 protein by MXL in the absence of baf. A1; however, when autophagy was inhibited by baf. A1, the inhibitory effect of MXL on NLRP3 was reversed (Figure 6A, B and S4). Further, the cells treated with MXL in the presence of baf. A1 showed a significant increase in the accumulation of LC3B-II, indicating stoppage of recycling of LC3B-II due to inhibition of autophagy (Figures 6A and B).

**Figure 6:**
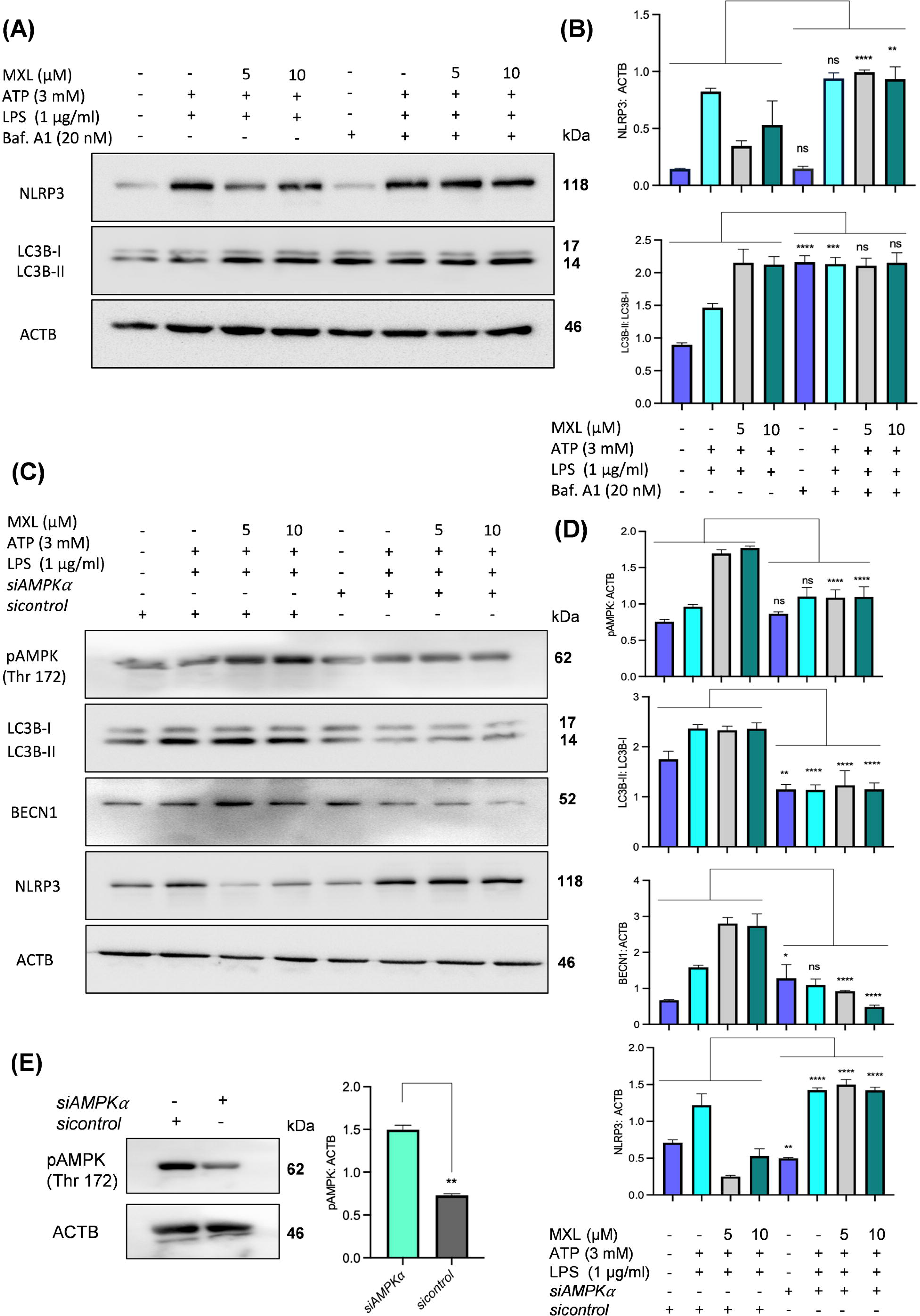
Pharmacological and genetic repression of autophagy limited the inhibitory effect of MXL on LPS and ATP induced NLRP3 inflammasome activation. Two approaches were utilized to evaluate MXL’s anti-NLRP3 inflammasome activity. Pharmacological inhibition of autophagy was achieved with bafilomycin A1, a late-stage autophagy inhibitor, and genetic repression was induced using siAMPK. The expression levels of autophagic proteins and NLRP3 inflammasome proteins were subsequently analyzed through western blotting. **(A)** Immunoblot analysis illustrating the protein levels of NLRP3 and LC3B. **(B)** Densitometric analysis of immunoblots revealed changes in protein expression of NLRP3: ACTB and LC3B-II: LC3B-I. **(C)** Immunoblot analysis of autophagic proteins (pAMPK, LC3B-II, BECN1) and NLRP3 protein. **(D)** Densitometric quantification of immunoblots showing expression levels of pAMPK: ACTB, LC3B-II: LC3B-I, BECN1: ACTB, NLRP3: ACTB. **(E)** Immunoblot analysis and densitometric quantification of pAMPK relative to ACTB. The data presented represent the mean ± SD of three independent experiments (n=3). Statistical analysis was performed using one-way ANOVA, followed by post-hoc Bonferroni test. A p-value <0.05 was considered statistically significant, with significance levels indicated as ****p < 0.0001, ***p < 0.001, **p< 0.01, *p< 0.05, and ns denoting not significant.

To confirm the MXL-induced autophagy by the AMPK pathway, we genetically knocked down AMPK by *siRNA* in J774A.1 cells followed by 1 h treatment with MXL. Results showed a significant reduction in pAMPK in the transfected control group, indicating successful knockdown (Figure 6E). The transfected cells treated with MXL showed no increase in pAMPK expression. However, the non-transfected group showed a significant induction of the AMPK pathway (Figures 6C and 6D). The expression of downstream proteins involved in autophagy regulation, such as BECN1 and LC3B-II, was reversed in transfected cells, which clearly indicated that MXL induces autophagy via the AMPK pathway (Figures 6C and D). Further, we examined whether MXL- induced inhibition of the NLRP3 inflammasome is related to MXL-induced autophagy via the AMPK pathway. To explore this relationship, the expression levels of proteins involved in NLRP3 inflammasome activation were evaluated. Results showed that AMPK knockdown reversed the effect of MXL on NLRP3 inflammasome inhibition, as evidenced by increased NLRP3 protein expression on western blotting (Figures 6C and D).

### 3.7 MXL inhibited the induction of NLRP3 inflammasome in two different in vivo NLRP3 inflammasome models

After evaluating the anti-NLRP3 inflammasome activity of MXL in the in vitro studies, we validated the efficacy of MXL in two different mouse models involving NLRP3 inflammasome-mediated inflammation. In both models, we used two doses of MXL (10 and 20 mg/kg). Firstly, the air mouse model was used to determine the effect of MXL on MSU-induced NLRP3 inflammasome. The lavage collected from air pouches was analyzed to evaluate the levels of different pro-inflammatory cytokines. The results showed that MXL treatment significantly reduced the NLRP3 inflammasome-mediated secretion of IL-1β and IL-18 as measured through ELISA. Colchicine, used in this study as a standard drug, showed similar activity to that of MXL (Figures 7A and B). The reduction in cytokine levels demonstrated the inhibitory effect of MXL on the NLRP3 inflammasome.

**Figure 7:**
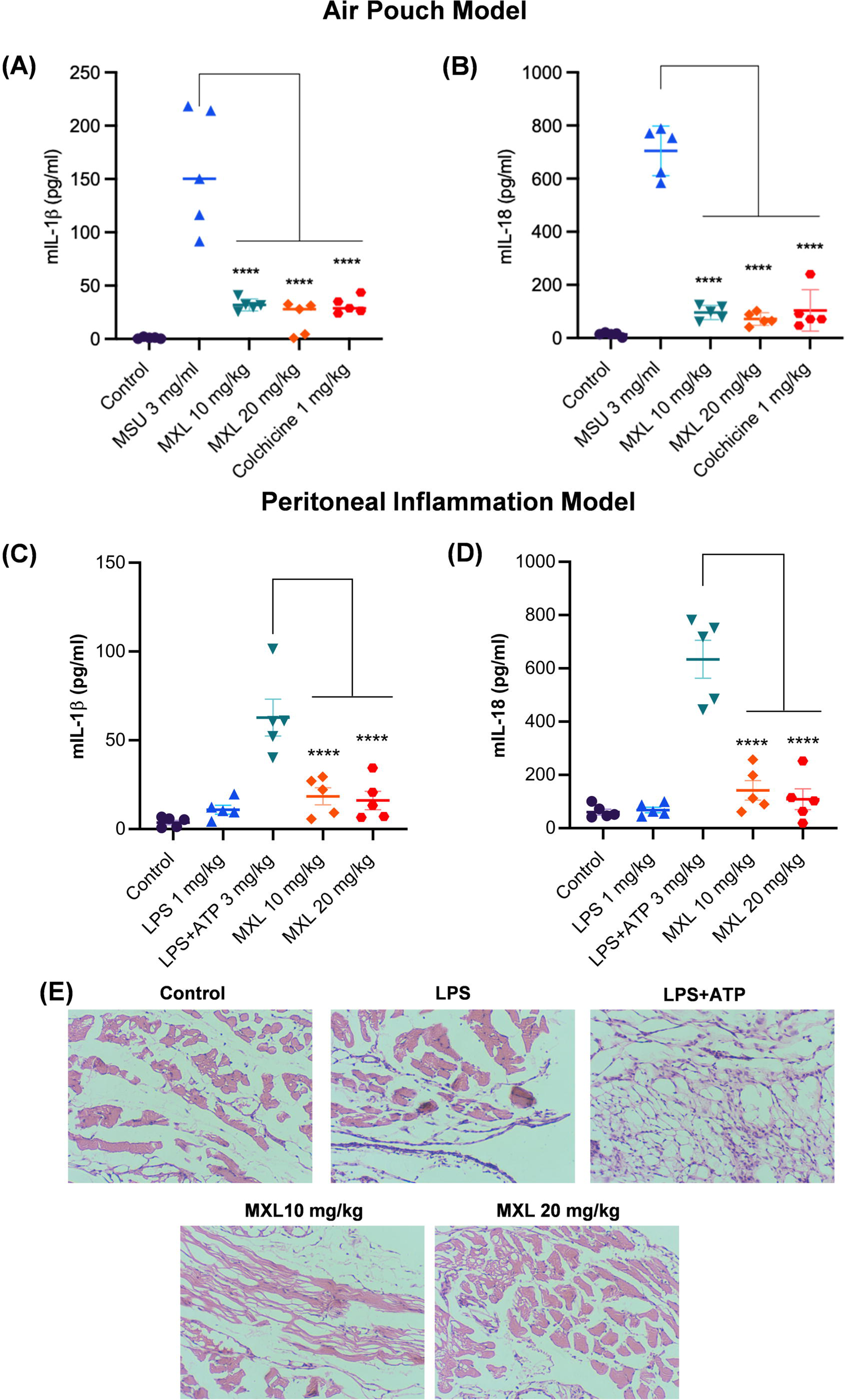
MXL inhibited the induction of NLRP3 inflammasome in two different in vivo NLRP3 inflammasome models. Female Balb/c mice were randomly assigned into five groups, each consisting of five animals (n=5). After administering the drug treatment, monosodium urate (MSU) crystals were injected into the induced air pouch for a duration of 6 hours. The collected lavage fluid was subsequently analyzed for the presence of NLRP3 inflammasome-mediated pro-inflammatory cytokines, IL-1β, and IL-18. **(A)** The graph illustrates IL-1β levels, and **(B)** shows IL-18 levels in the collected lavage fluid. In the LPS/ATP-induced peritoneal inflammation model, drug treatment was administered before the administration of LPS and ATP. The peritoneal lavage fluid was then evaluated for levels of pro-inflammatory cytokines. **(C)** The graph represents IL-1β levels. **(D)** The graph represents IL-18 levels in the peritoneal lavage fluid. **(E)** Representative microscopy images illustrate the histological analysis of peritoneal tissue in the following groups: control, LPS, LPS+ATP, MXL 10 mg/kg, and MXL 20 mg/kg treated mice. Statistical analysis involved one-way ANOVA followed by a post-hoc Bonferroni test. A p-value <0.05 was deemed statistically significant, with levels of significance annotated as ****p < 0.0001, ***p < 0.001, **p< 0.01, *p< 0.05, and ns indicating non-significant.

Further, we utilized the LPS/ATP-induced peritoneal inflammation model to investigate the in vivo effects of MXL, and the levels of pro-inflammatory cytokines were analyzed in peritoneal lavages using ELISA. The findings demonstrated that MXL treatment significantly inhibited the NLRP3 inflammasome-mediated release of IL-1β and IL-18. These data clearly demonstrated the inhibitory effect of MXL on LPS-and ATP-induced pro-inflammatory cytokines in the peritoneal inflammation model (Figures 7C and D). The attenuation of LPS and ATP-induced inflammation by MXL treatment was demonstrated through histological analysis of peritoneal tissue. In mice treated with LPS and ATP, histological examination revealed disorganized tissue morphology, indistinct boundaries between the dermal and hypodermal layers, and notable cellular ballooning (Figure 6E). However, these pathological features were clearly prevented in mice pretreated with MXL (Figure 7E).

## 4. Discussion

The NLRP3 inflammasome is a crucial element of the innate immune system, and its activation can be triggered by a broad spectrum of pathogen-associated molecular patterns (PAMPs) and damage-associated molecular patterns (DAMPs) (12–13). Nevertheless, its dysregulation contributes to the pathogenesis of numerous inflammatory diseases, such as Inflammatory bowel disease (IBD), atherosclerosis, and Alzheimer’s disease (AD), etc., through the secretion of pro-inflammatory cytokines IL-1β and IL-18 (14–15). Therefore, multiple approaches, including the repurposing of drugs, are being used to identify inhibitors of the NLRP3 inflammasome for the treatment of inflammatory diseases. In this study, we screened some of the US FDA drugs to identify inhibitors of the NLRP3 inflammasome. We identified two drugs, including fluvoxamine and minoxidil, during screening. However, this study is focused on minoxidil, as we have already published our work on fluvoxamine (11). Minoxidil topical solution is used for the treatment of androgenetic alopecia, while it is orally used as an anti-hypertensive drug. This is the first report indicating the anti-NLRP3 inflammasome activity of minoxidil.

Using J774A.1 murine macrophages stimulated with LPS and ATP, we observed that MXL significantly inhibited the NLRP3 inflammasome-mediated secretion of IL-1β and IL-18. The anti-NLRP3 inflammasome activity of MXL was also confirmed in non-cancerous BMDM cells, indicating a generalized pharmacological effect of MXL against NLRP3 inflammasome. Furthermore, we evaluated the impact of MXL on key proteins involved in activating the NLRP3 inflammasome and observed a significant reduction in the expression of NLRP3, CASP1, IL-1β, along with decreased ASC oligomerization. Additionally, we aimed to investigate the underlying mechanism contributing to the anti-NLRP3 inflammasome activity of MXL. The involvement of dysregulated NLRP3 in multiple inflammatory diseases requires its precise regulation under physiological conditions to ensure adequate immune protection while preventing damage to host tissue. In this context, autophagy acts as the primary regulatory mechanism to mitigate the over-activation of the NLRP3 inflammasome in various pathological conditions (16–18). Autophagy mediates the lysosomal degradation of damaged proteins, organelles, NLRP3 inflammasome activators, inflammasome-associated components, and cytokines. Numerous studies have demonstrated that pharmacological upregulation of autophagy can alleviate the activation of the NLRP3 inflammasome (19–26).

Minoxidil has been reported to induce autophagy in cells of different origins (27–28). Therefore, we investigated the involvement of autophagy in MXL-mediated inhibition of NLRP3 inflammasome. We found that under inflammatory conditions, MXL treatment led to upregulation of the autophagic proteins pAMPK (Thr172), BECN1, and LC3B-II, along with decreased expression of pMTOR (Ser2448), SQSTM1, and 4E-BP1. These findings suggested that MXL exerted its anti-NLRP3 inflammasome effect by inducing autophagy under inflammatory conditions.

To confirm the involvement of autophagy in the anti-inflammatory effects of MXL, we inhibited autophagy by using pharmacological and genetic approaches. The pharmacological inhibition of autophagy reversed the anti-NLRP3 inflammasome activity of MXL, confirming the involvement of autophagy. While *siAMPK*-mediated knockdown also exerted similar effects, it additionally confirmed the involvement of AMPK-mediated induction of autophagy by MXL in regulating NLRP3 inflammasome activity. These findings demonstrated a novel MXL-mediated interplay between the NLRP3 inflammasome and autophagy. Our results also corroborate the data on other AMPK activators like metformin, AICAR, and A-769662 in the modulation of NLRP3 inflammasome activity by inducing autophagy (29–32).

Our findings about the anti-NLRP3 inflammasome activity of MXL present a paradoxical picture as MXL exerts its vasodilator activity by causing of potassium efflux through K(ATP) channels (33). Potassium efflux is a canonical mechanism for induction of the NLRP3 inflammasome; therefore, it presents an opportunity to further study the paradoxical role of MXL in the context of potassium efflux and NLRP3 inflammasome activation.

After confirming the molecular mechanism of MXL against NRLP3 inflammasome, we validated the anti-NLRP3 inflammasome activity in two well-established NLRP3-mediated inflammation models: monosodium urate (MSU) induced air pouch model and LPS/ATP induced peritoneal inflammation model. Peritoneal inflammation was induced using the established LPS-priming and ATP-stimulation model, which activates NLRP3 inflammasome signaling to drive IL-1β and IL-18 maturation and release in vivo. Similarly, the monosodium urate (MSU) induced air pouch model involved subcutaneous injection of MSU crystals to trigger neutrophil influx and pro-inflammatory cytokines production (34–37). These models were chosen for their relevance in mimicking inflammatory responses mediated by the NLRP3 inflammasome. The findings revealed that MXL exerts anti-NLRP3 inflammasome activity in both models. In the MSU-induced air pouch model, MXL treatment significantly reduced IL-1β and IL-18 levels, indicating potent inhibition of NLRP3 inflammasome activation. Similarly, in the LPS and ATP-induced peritoneal inflammation model, MXL administration markedly attenuated the secretion of IL-1β and IL-18, further corroborating its efficacy in suppressing NLRP3 inflammasome activity.

This investigation revealed MXL as a potent inhibitor of the NLRP3 inflammasome. We propose to further explore its potential as a therapeutic agent for conditions in which overactivation of the NLRP3 inflammasome exacerbates inflammation.

## 5. Conclusion

MXL was investigated for its role in inhibiting the NLRP3 inflammasome. Our studies demonstrated that MXL promotes autophagy via activation of the AMPK pathway, which modulates the NLRP3 inflammasome activity. Downregulation of AMPK via siRNA and pharmacological inhibition of autophagy led to a marked upregulation of NLRP3, underscoring the role of autophagy in regulating NLRP3 via MXL. Furthermore, the anti-NLRP3 activity of MXL was validated in two distinct in vivo models of NLRP3 inflammasome-mediated inflammation. Our findings warrant further studies to develop minoxidil as a possible therapeutic agent for NLRP3-mediated inflammatory diseases.

### Limitations of the study

This study indicates that MXL modulates activity of the NLRP3 inflammasome through AMPK-mediated autophagy. However, these findings do not rule out direct interaction between MXL and other proteins in the NLRP3 inflammasome complex. Apart from that, validation of the molecular mechanism of action of MXL and its pharmacokinetic profiling to determine the optimal in vivo dose for modulating NLRP3 inflammasome-mediated inflammation will further strengthen the findings of this study. These studies will be taken up as part of future investigations to expand knowledge of the modulation of NLRP3 inflammasome activity by MXL.

## Statement and Declarations

## Competing interests

The authors declare that they have no competing interests.

## Acknowledgments

We are thankful to the Council of Scientific and Industrial Research (CSIR), India, for providing research fellowships to Sukhleen Kaur, Mehboob Ali, and Lakhvinder Singh during the course of this work. We are also thankful to the Department of Science and Technology (DST), India, for the research fellowship to Ziya Tufail. We are also thankful to CSIR-Indian Institute of Integrative Medicine, Jammu, for financial support for this work under the project MLP-6002 (WP-4) and partial financial support through the project MLP-002622.

## Ethics Statement

This study used mice. All necessary approvals for the use of animals were obtained by the authors from the animal ethics committee of the CSIR-Indian Institute of Integrative Medicine, Jammu. The authors also declare that all data shown in this manuscript are accurate to the best of their knowledge.

## Authors’ contributions

SK and MA performed in vitro and in vivo experiments in this study. AS, ZT and LS contributed to some of the in vitro experiments and helped in the analysis of data. ZA contributed to the study design and provided institutional support. SK and AK designed the study, analyzed the data, and wrote the manuscript.

## Notes

### Competing Interest Statement

The authors have declared no competing interest.

### Summary of Updates

We revised the manuscript to include newly generated data, including an analysis of minoxidil's anti-NLRP3 activity in bone marrow-derived macrophages (BMDMs). In addition, we tested the effect of minoxidil on cell death under inflammatory conditions. The new data have been added to the figures as Figures 2A, B, C, D, F and Figures 3E, F and G. We have revised the abstract, results, and discussion sections as well. The additional data were generated to two more authors. Therefore, their names have been included in the revised manuscript.

