## Supplementary Information File for "Minoxidil hydrochloride impedes NLRP3 inflammasome activation via upregulation of AMPK-mediated autophagy"

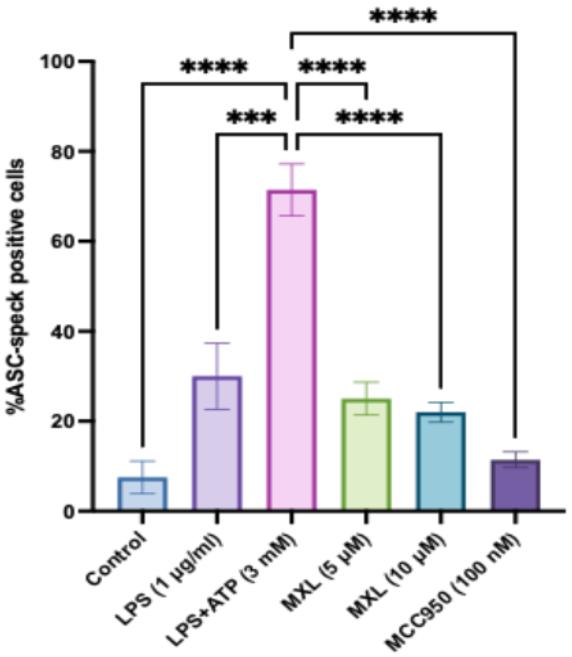


**Figure S1**: MXL suppresses LPS+ATP-induced ASC speck formation. Cells were treated with LPS (1 μg/mL) and ATP (3 mM) to induce ASC speck formation and subsequently treated with MXL (5 or 10 μM) or the NLRP3 inflammasome inhibitor MCC950 (100 nM). ASC speck formation was quantified as the percentage of ASC-speck–positive cells. LPS+ATP treatment markedly increased the proportion of ASC-speck-positive cells compared with the control group, whereas treatment with MXL (5 and 10 μM) significantly reduced LPS+ATP-induced ASC speck formation. Data are presented as mean ± SEM. Statistical analysis was performed using one-way ANOVA followed by Bonferroni's post hoc multiple-comparisons test. *** $p < 0.001$ and **** $p < 0.0001$.
